# Time series analysis in a maize landrace reveals rapid fixation of beneficial alleles

**DOI:** 10.64898/2026.03.25.713886

**Authors:** Margarita Takou, Michelle Terán-Pineda, Sofia da Silva, Chris-Carolin Schön, Markus G Stetter

## Abstract

Identifying loci in the genome that allow a population to respond to selection pressure is essential to understand evolution and improve crops. Temporally consecutive generations under selection offer the opportunity to identify signatures of selection. Maize, as one of the most important crops worldwide is rich in genetic diversity and a model for breeding advances. Therefore, it is an ideal system to study genetic changes in response to selection. Here, we study the genetic changes in two replicates of a selection experiment in a European maize landrace, which showed rapid trait improvement over three cycles of selection. We identified an increase in genetic divergence across successive doubled-haploid populations derived from each selection cycle, consistent with the effect of strong directional selection. The genetic divergence observed between the replicates was greater than that between generations. In addition to the genome-wide signal, we identified multiple candidate loci under selection through temporal F_ST_ outlier analysis comparing the original landrace population to subsequent cycles. These loci showed a significant overlap with genomic regions, controlling intentionally selected traits and other traits. The significant overlap of selected loci between the two replicates shows the importance of major loci in response to directional selection, while the large number of non-overlapping loci demonstrates the polygenic response. Our work shows that the temporal dimension in plant breeding time-series enables the identification of candidate loci under selection and the genome-wide dynamics of change in response to selection.

## Introduction

The evolutionary forces shaping the levels of genetic diversity within populations are genetic drift and selection (Ellegren and Galtier 2016; Leffler *et al*. 2012). Both forces co-occur and shape the changing pattern of diversity along the genome as time progresses in finite populations. While genetic drift alone can lead to substantial differentiation between populations (Charlesworth 2009), directional selection amplifies differentiation, particularly at the genetic loci under selection. In sufficiently large populations, selection will dominate the population and decrease the level of genetic diversity around beneficial alleles due to linkage disequilibrium in the genome (Smith and Haigh 1974; Wright 1931; Castellano *et al*. 2018; Cvijović *et al*. 2018; Charlesworth *et al*. 1993, 1995). The rapid change in allele frequencies at selected loci and linked neutral loci over time provides the opportunity to identify targets of selection (Barghi *et al*. 2020). How the overall diversity and selected loci shape the response to selection is an essential question in evolutionary biology, as well as for plant and animal breeding.

Time series experiments are powerful tools for studying the evolution of traits and how genetic diversity changes over time (Lai and Schlötterer 2022; Wu *et al*. 2025). For example, Oakley *et al*. (2023) were able to identify genetic loci involved in a fitness-related genetic trade-off, enabling the adaptation of *Arabidopsis thaliana* populations to local environments. Despite a few recently emerging studies in plant populations (Wu *et al*. 2025), the use of time series and experimental evolution has mostly been limited to microbes and model organisms such as *Drosophila melanogaster* and mice (Schlötterer 2023; Chan et al 2012). An established system of directional selection is plant breeding, where consecutive generations are selected for specific traits (Lopez-Reynoso & Hallauer 1998; Hallauer *et al*. 2004; Duvick *et al*. 2004; Moose *et al*. 2004). Hence, sampling and genotyping consecutive generations of plant breeding cycles provide the opportunity to identify selected loci and their differentiation over time.

In plant breeding, genome-based selection is used to accelerate progress and maximize selective gain (Heffner *et al*. 2010). The method has revolutionized breeding programs, as it enables the identification of superior individuals based on their genomic information. Strong directional selection can lead to rapid genetic diversity loss, increased population structure, and increased linkage disequilibrium (Dreisigacker *et al*. 2023; Arguello-Blanco and Sneller 2023). However, causal genetic regions under selection are usually not identified. Consecutive plant breeding cycles, in combination with population genetic inference, can give insights into selection progress and help to identify adaptive alleles that contribute to the responses to selection.

As one of the most important crops worldwide, maize has been at the forefront of the development of advanced breeding technologies for decades. The high genetic diversity available in landraces and its domestication history have made maize also a model crop for evolutionary biology (Yang *et al*. 2023; Hufford *et al*. 2012). The introduction of maize to Europe has shaped a distinct set of landraces that enabled the successful establishment of the crop across the European continent (Tenaillon and Charcosset 2011; Takou *et al*. 2025). Doubled-haploid (DH) lines derived from these landraces have successfully been used to identify genetic loci controlling important traits including flowering time and early plant development (Mayer *et al*. 2020). Hence, these lines are highly useful to understand functional diversity and selection.

In a selection experiment starting from 402 DH lines of the landrace Petkuser Ferdinand Rot, Polzer *et al*. (2025) selected for early plant development (plant height measured at stages V4 and V6) and imposed stabilizing selection on final plant height by selecting the top 20 lines to create two sets of founder lines. Each set of lines was mated in a diallel crossing scheme. Selection was applied in two subsequent cycles based on the genomic breeding values of heterozygous individuals. DH populations were produced in each cycle and used as a proxy for the selection units. Selection was successful for traits under directional selection. The largest phenotypic response to selection was observed in the first cycle and decreased in later cycles. The two replicates differed in selection response. While the successful phenotypic response has been revealed through a common garden experiment of the DH lines from consecutive cycles, the genetic loci responsible for the outcome has remained unknown.

Here, we study the changes in the genetic composition of consecutive populations in experimental cycles across two replicates. We find that genetic differentiation increases more between replicates than between cycles of the same replicate and that highly beneficial genomic loci fixed early in the experiment, explaining the high selection gain in the first generation. Selection scans across the genome identify functionally important variation that contributes to the traits under selection, but also loci that have potentially been under ‘natural selection’ during the experiment. Our results shed light on the short-term genetic changes during directional selection in maize.

## Materials and Methods

### Rapid cycling selection experiment

We study the genomic changes during directional selection in a maize breeding experiment. The initial rapid cycling experiment has been described Polzer *et al*. (2025) and was developed based on material from Hölker *et al*. (2019) and Mayer *et al*. (2022). The initial goal of the experiment was to investigate the potential of rapid cycling genomic selection for pre-breeding of maize landraces. The genomic prediction model for selection was trained on 899 flint doubled-haploid (DH) lines from the European maize landraces Petkuser Ferdinand Rot (PE, Germany), Kemater Landmais Gelb (KE, Austria) and Lalin (LL, Spain). Twenty-five traits of interest were assessed across 11 environments within Europe. The selection criterion combined directional selection for early plant height (stages V4 and V6) and stabilizing selection for final plant height. Three cycles of genomic selection and interbreeding were performed in two replicates (R1 and R2). A detailed description of the scheme can be found in Polzer *et al*. (2025). In short, from the base population (C0) of 402 DH lines derived from landrace PE, top-ranking DH lines were selected based on breeding values and partitioned into founder lines for R1 (N = 10) and R2 (N = 8). These founders were interbred in a diallel scheme, and the resulting offspring were selfed once. Genomic selection was applied, and selected plants were crossed maximizing pairwise modified Rogers’ distance (Wright 1978). This process of selection and paired crossing was repeated in cycle 2. DH lines were extracted from each cycle and replicate for evaluation (Figure 1a).

**Figure 1.**
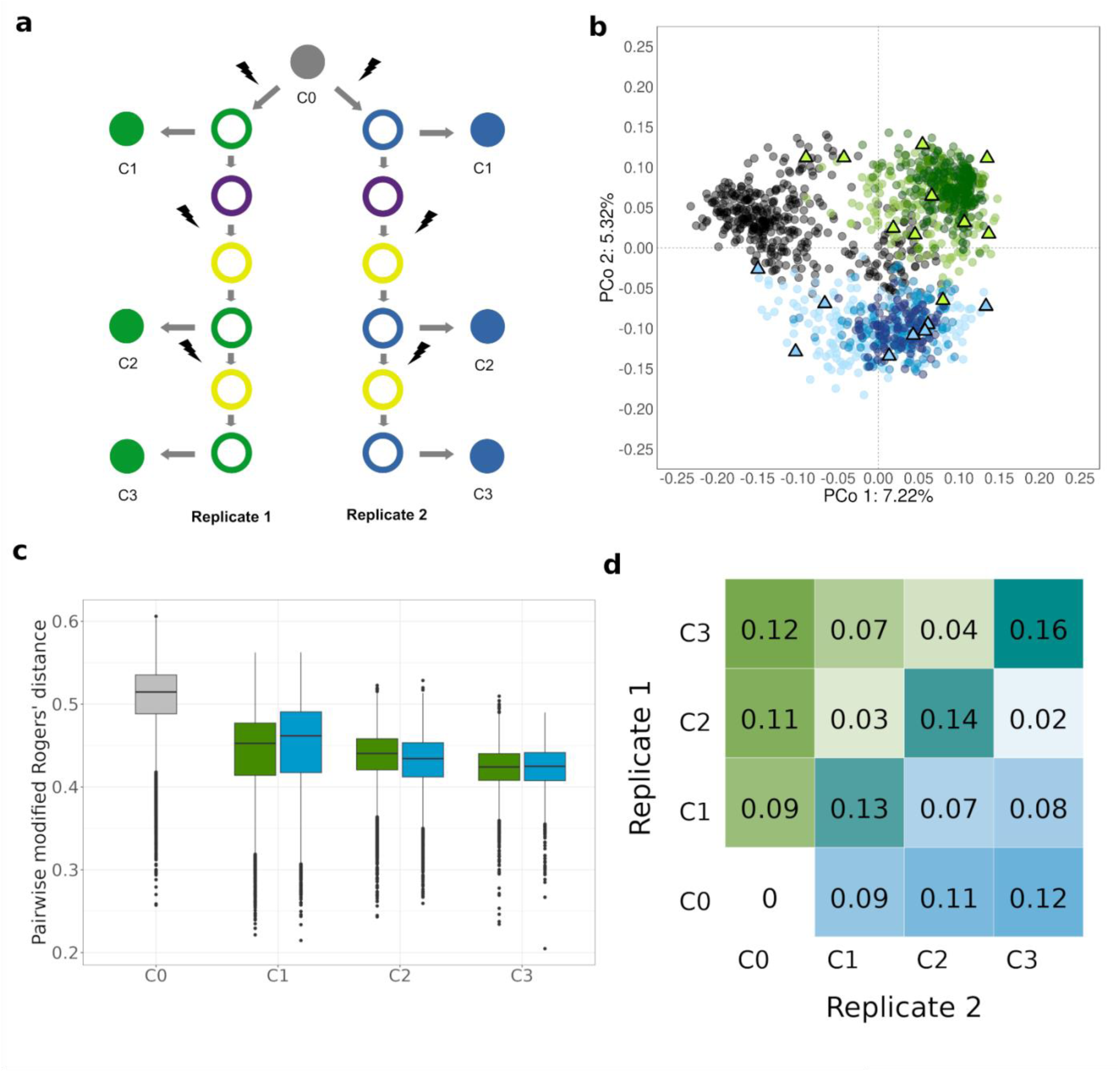
Genetic differentiation of the seven populations in the selection experiment. **a.** Schematic representation of the selection experiment. Starting with an initial population of Petkuser landrace DH lines (C0; grey filled circle), three cycles of genomic selection (indicated by the black bolts) were applied to each replicate, R1 (green) and R2 (blue). Purple and yellow circles represent selfing and interbreeding steps, respectively; filled circles depict the produced DH populations that were used in this analysis. b. Principal Coordinate Analysis (PCoA) of DH lines based on pairwise modified Rogers’ distances. Grey points represent the base population (C0). C1 through C3 are depicted in green (R1) and blue (R2), with darker shades indicating later cycles. Triangles denote founder lines. c. Modified Rogers’ distances (MRD) calculated pairwise among DH lines within each population (C0-C3) for both replicates. d. Hudson’s genome-wide *F_ST_* values within and between cycles and replicates. The upper triangle displays *F_ST_* values for R1, and the lower triangle for R2; the diagonal shows *F_ST_* values comparing the two replicates.

Our study investigates seven populations, totaling 1,258 DH lines. These include the shared base population (C0) of 402 lines and the subsequent three cycles (C1 to C3) in each replicate. R1 consists of 168 lines each for C1 and C2, and 114 for C3. The second replicate R2 comprises 159, 160, and 87 lines for cycles 1 through 3, respectively (Supplementary Table S1). Our dataset includes 11,160 SNP markers, which covered the whole genome. Marker distribution was visualized in CMplot (v4.5.1; Supplemental Figure S1).

### Population structure and genetic diversity

We analyzed the population structure across all cycles and replicates with a principal coordinate analysis (PCoA; Gower 1966) based on pairwise modified Rogers’ distances (MRD; Wright (1978)) between DH lines. Analyses were performed with the R (v4.4.3) package ape (v5.8.1; Paradis *et al*. (2019)). Genome-wide genetic differentiation was calculated as the average Hudson’s fixation index (*F_ST_*) across all SNPs. These pairwise calculations were performed for populations within each replicate and for corresponding cycles between replicates using the R package KRIS (v1.1.6; Chaichoompu *et al*. (2018)). To assess genetic diversity within each cycle and replicate, we analyzed each population separately, using the 9,668 SNPs that were polymorphic in C0. We estimated the proportion of polymorphic markers (P), and gene diversity (*H*; Nei (1973)) for each population (Mayer et al., 2020). We compared distributions of population structure, and genetic diversity measures with the non-parametric, two-sample Kolmogorov-Smirnov (KS) test in R v4.1.2 to test for significant differences between the populations (Marsaglia et al 2003).

### Genome scans for loci under selection

To identify regions in the genome that were under selection, we compared genetic differentiation per SNP between each selection generation (C1 to C3) to the training population (C0). We calculated pairwise *F_ST_* along the genome, using Hudson’s *F_ST_* with the scikit allel package v1.3.7 (Miles *et al*. 2023) for python v3.9. Significant signals of selection were identified, by permuting the identity of the individuals 1,000 times. The cut-off was the 99^th^ percentile permuted *F_ST_* values. We further evaluated the robustness of the selection signal through the comparison of the recall of selection signals across selection cycles. We overlapped a 20 kb genomic region around each significant SNP (10kb up- and downstream) with regions of genome-wide association study (GWAS) results for multiple important phenotypes. These associations were found in the initial Petkuser DH population (Mayer *et al*. 2020). The overlap between the selection signals and GWAS results was tested by a hypergeometric test at a significance level of 0.05 (Casella and Berger, 2004; Rivals et al 2007). In addition, we performed a gene ontology (GO) enrichment analysis for genes within the selected regions, using the topGO package with the ‘elim’ algorithm and a Fisher’s exact test at a significance level of p <0.05 (Alexa *et al*. 2010). We also tested GO enrichment specific to each replicate by excluding common significant genes. All data were plotted using the ggplot2 library in R v4.1.2 (Wickham 2016).

## Results

### Genetic divergence between replicates increases over time

Directional selection is expected to increase genetic differentiation to the initial population over time. To assess the impact of selection on the genome-wide differentiation of the experimental populations, we investigated changes in population structure and genetic diversity. A principal coordinate analysis (PCoA) of the seven populations using 11k SNP markers revealed that genetic structure was primarily defined by selection cycle and replicate (Figure 1b). The first principal coordinate accounted for 7.22% of the total variance and captured the progressive divergence of selection cycles (C1–C3) from the base population C0 in both replicates. Replicate 1 was overall more diverged from C0 than R2. The second principal coordinate explained 5.32% of the variance, reflecting the clear divergence between replicates R1 and R2. Furthermore, the increased clustering density observed from C1 to C3 suggested a progressive genetic separation as selection cycles advanced.

The genome-wide genetic distances between DH lines within each cycle declined progressively over time, a trend consistently observed in both replicates (Figure 1c). The distribution of the modified Rogers’ distances (MRD) in C1 had median values of 0.45 and 0.46 for R1 and R2, respectively. The MRD decreased over time, with median values of 0.44 and 0.42 for C2 and C3 in R1, respectively, and 0.43 for C2 and C3 in R2, respectively. The standard deviations of the MRD values also decreased within both replicates across cycles. For example, within R1, the standard deviations changed over time from 0.05 in C1 to 0.03 in C2 and C3, a pattern which corroborates the PCoA clustering (Figure 1b). MRD distributions differed significantly between replicates at each corresponding cycle, as indicated by the KS test results of p < 2.2e-16 for the comparisons within C1 and C2, and p = 0.035 for the comparisons within C3 (Figure 1c). Thus, the two replicates diverged more than they changed over time within each replicate.

While genetic distances within each cycle decreased, genetic differentiation between cycles within each replicate did not increase over time. In R1, genetic differentiation was maximal between C0 and C1 (Hudson’s genome-wide mean *F_ST_* of 0.09; Figure 1d), with lower mean *F_ST_* values observed between C1 and C2 (0.03) and between C2 and C3 (0.04). R2 exhibited a similar trend, with the most substantial shift occurring between C0 to C1 (mean *F_ST_* = 0.09); however, differentiation remained relatively high between C1 and C2 (0.07) before declining to 0.02 between C2 and C3. Across all cycles, mean *F_ST_* values between replicates exceeded those within replicates, corroborating the PCoA results. Differentiation between R1 and R2 was 0.13 at C1, rising to 0.16 in C3. These results show that selection and genetic drift shaped the consecutive cycles of the experiment, consistent with significant changes in phenotypes over time (Polzer *et al*. 2025).

### Genetic diversity decreases during the selection experiment

While the increase in genetic differentiation over time within each replicate was relatively low and stable, the variation between lines declined. Therefore, we investigated changes in the genetic diversity between generations in more detail. The proportion of polymorphic loci decreased by approximately 35% in both replicates in the first selection cycle (Figure 2). There was no decrease between the founder lines and C1. Given that no selection has been applied between the founders and C1, this confirms that divergence in generations without selection was rather low in the experiment. The genetic diversity decreased within both replicates during the following cycles, C2 and C3.

**Figure 2.**
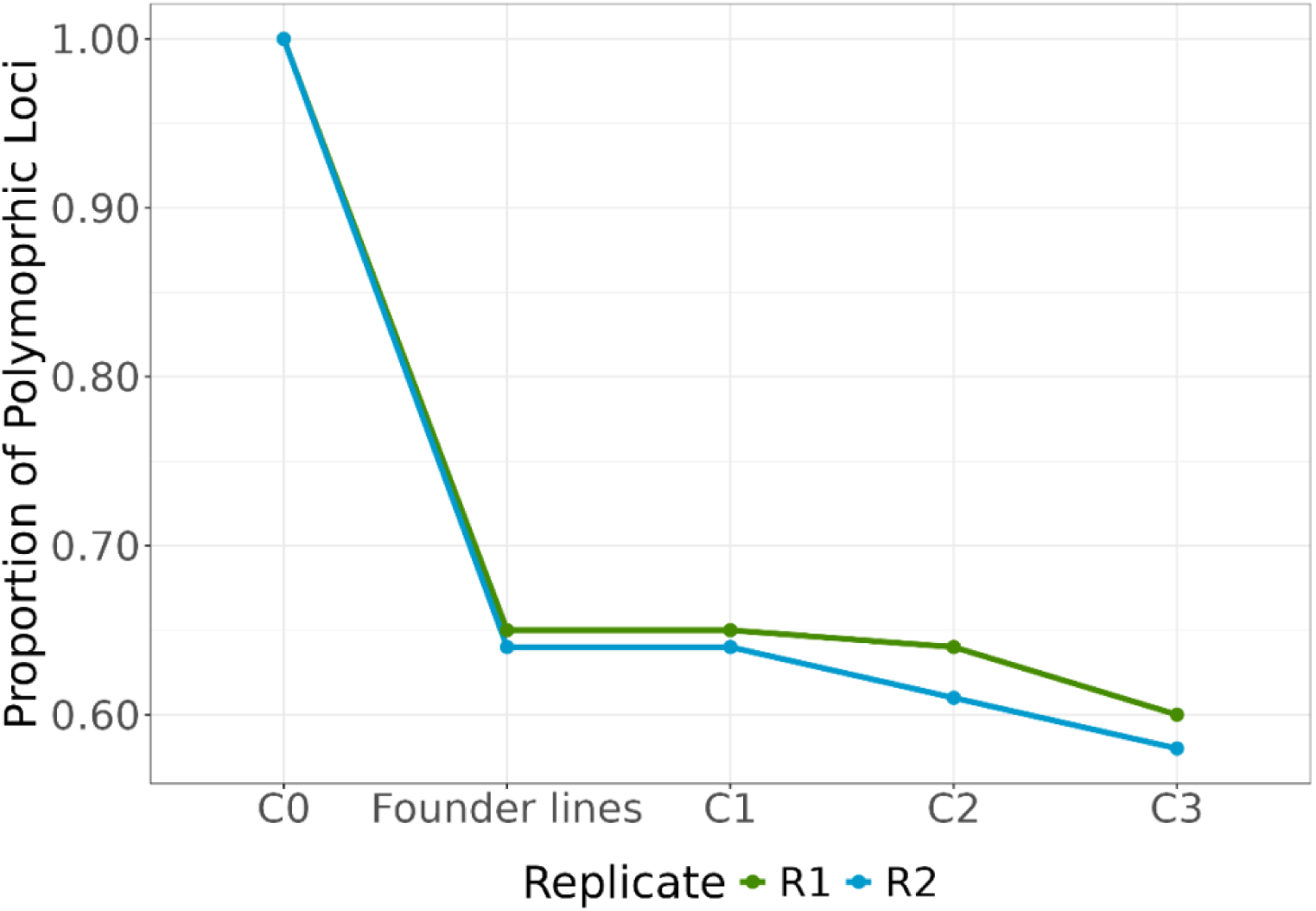
Proportion of polymorphic SNPs compared to the initial population. Segregating sites out of 9,668 polymorphic SNPs that were polymorphic in population C0.

The gene diversity (*H*) across all loci was highest in C0, with a median of 0.33 and a mean value of 0.30 (Figure S2). The *H* in the 18 founder lines declined to medians of 0.32 in R1 and 0.22 in R2 (p < 2.2e-16, KS test). From C1 to C3, median *H* further decreased to 0.19 in R1 and R2. Across both replicates, *H* in cycles C1–C3 were significantly different from C0 (KS test, p<8.312e-11). We detected a decrease in genetic diversity at each cycle of the experiment and observed a significant difference in genetic diversity levels between replicates, suggesting a polygenic nature of differentiation.

### Parallel selection drives the frequency increase of different loci between replicates

We investigated signatures of selection across the genome over the consecutive cycles of the experiment. As loci under strong directional selection should change genetic differentiation between generations, preferably at selected loci, we estimated the *F_ST_* along the genome between C0 and each of the subsequent cycles (C1 to C3). To find consistent and robust signatures of selection, we considered *F_ST_* values that were outliers (permutation based p < 0.05) in at least two of the pairwise comparisons to C0 (Figure 3a; Figure S3). This should decrease the risk of false positives caused by fluctuations in allele frequencies due to genetic drift between populations. Out of the 11,160 genetic markers tested, we identified 297 and 232 outliers for R1 and R2, respectively. The outliers were distributed across the genome, with few larger clusters on Chromosome 1, Chromosome 4, and Chromosome 10 in R1 (Figure 3a). On Chromosome 1, 27 outliers were shared in all four comparisons. The outliers in R2 were also distributed across the genome (Figure S4), with 34 loci significantly overlapping between the two replicates (hypergeometric test p-value of <2.2e-16; Figure 3b). This observation aligns with the interpretation that the phenotypes under selection have a polygenic basis with outlier loci that might represent the largest fitness effects.

**Figure 3.**
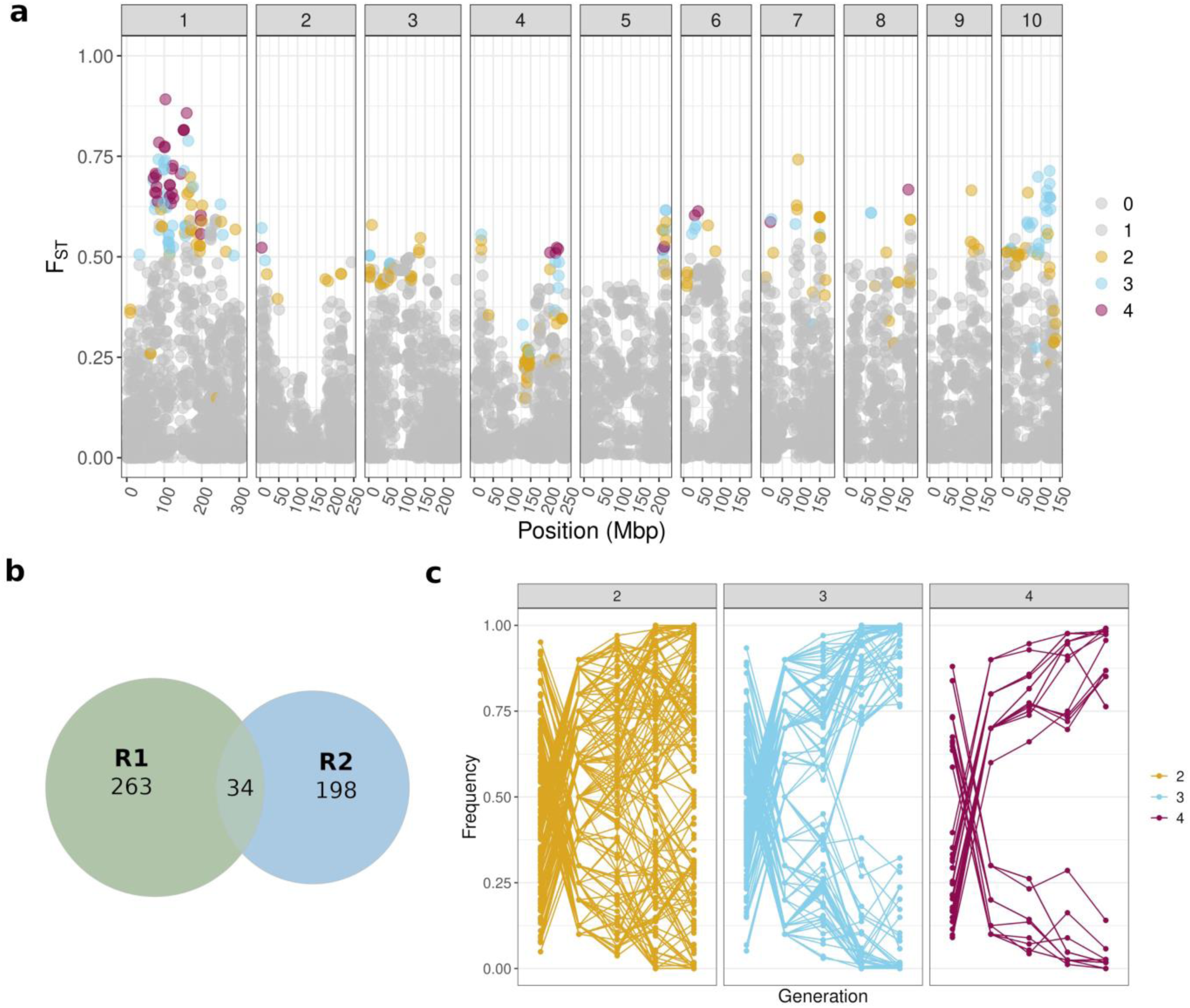
Signals of selection during the selection experiment. **a.** The *F_ST_* values between C0 and C3 indicate changes in R1. The colored dots in both plots indicate in how many of the comparisons between C0 and founder lines, C1, C2, and C3, the locus was an outlier for *F_ST_*, while each chromosome is shown in its own facet. b. Significant overlap (p < 2.2e-16) of 34 outlier loci between R1 and R2. c. Within R1, the outlier loci are determined early in the selection cycle based on allele frequency changes.

Most of these loci were possibly selected early in the experiment, as their allele frequencies strongly increased from C0 to C1 in both replicates (Figure 3c). Outlier loci found in all four comparisons in R1 had, on average, a larger genetic differentiation (mean *F_ST_* = 0.52) from C0 to C1 than loci that were outliers in only three comparisons (mean *F_ST_* = 0.37). In R2, the mean *F_ST_* was 0.60 and 0.29 for outliers in all cycles and three comparisons, respectively. Within R1 45% of the outliers in two cycles had an allele frequency above 0.80 (or below 0.20) in C1, while this percentage rose to 60% of the outliers in three cycles and 100% of the outliers in all comparisons to C0. Results were similar in R2; 36.40% and 100% of the outliers in two comparisons and four comparisons to C0, respectively, were close to fixation for either allele in C1. However, the percentage was 8% of the outliers in three comparisons. Thus, a substantial proportion of the selected loci comes close to fixation in the first selection step.

### Phenotypes and biological functions are repeatedly selected

To understand the potential role of the candidate loci for selection, we examined overlaps of 20 kb regions (10 kb upstream and downstream of significant markers) with previously identified QTL, controlling eight growth-related traits in European maize, including the PE landrace (Mayer *et al*. 2020). We identified a significant overlap of the outlier loci (p = 6.065e-07) in R1 with seven QTL (Figure 4). Four of those QTL were located on Chromosome 1 of the maize genome. These were associated with the traits early vigor at V4 and V6 and final plant height, on which the population had been selected. In R2, one of four significant (p = 3.31e-27) overlaps with a QTL was related to the selected phenotype ‘plant height at the V4 stage’. Both replicates shared the same selected region on Chromosome 5 that overlapped with a QTL controlling lodging, a trait not included in the selection criterion, indicating that the trait had been under selection in the field.

**Figure 4.**
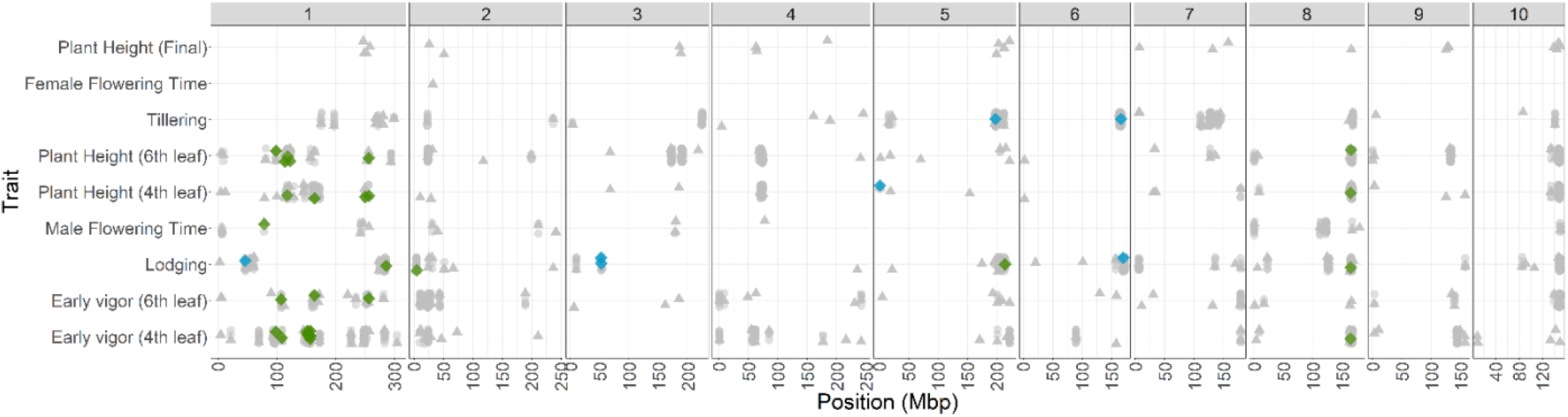
Biological meaning of sites with high genetic differentiation. The outlier *F_ST_* values overlap with known regions controlling phenotypes in European landraces. The green and blue diamonds indicate the loci that were outliers in multiple generations and overlap with the GWAS windows for R1 and R2, respectively. The grey points are overlaps of outliers in one generation only and the triangles indicate the GWAS window starting point.

Functional signatures provided a characterization of the identified selection signals. We extracted 371and 294 genes within the identified windows in R1 and R2, respectively, comprising shared and non-shared regions between the two replicates. Gene Ontology (GO) enrichment analysis identified 62 enriched GO terms in R1 (Table S2) and 79 in R2 (Table S3). Among those, six GO terms were significantly enriched in both replicates and included functions such as ‘acquisition of seed longevity’, ‘negative regulation of reproductive process’, and ‘indeterminate inflorescence morphogenesis’.

The overlap of categories between replicates was significant (p < 2.2e-16) attributable to shared selected markers between the two replicates. The removal of these shared markers from the analysis, eliminated all common enriched GO terms, suggesting parallel selection on major alleles in both replicates. Investigation into the functional level of parallel selection revealed that genes unique to each replicate favored diverse biological functions. The top enriched GO terms identified in R1 featured ‘dicarboxylic acid biosynthetic process’ (p = 0.00059), ‘indeterminate inflorescence morphogenesis’ (p = 0.00097), ‘polyprenol biosynthetic process’ (p = 0.00394), and ‘PSII associated light-harvesting complex II catabolic processes’ (p = 0.00426). Significantly enriched GO terms in R1 included ‘sterol metabolic process’ (p = 0.0026), ‘post-embryonic development’ (p = 0.0035), ‘hexose phosphate transport’ (p = 0.0069), and ‘regulation of TOR signaling’ (p = 0.0069). Therefore, there is not only a significant overlap between selected loci and selected traits, but also evidence for selection on functional units.

## Discussion

Tracking the effect of directional selection on different levels of genetic diversity over time can help to understand the forces that shape diversity and enable efficient plant breeding. Here, we show that after a strong initial decrease in genetic diversity, the loss of diversity decreased only moderately over time (Figure 2). The observed plateau in genetic differentiation and genetic diversity between cycles might explain the relatively small phenotypic progress observed after the first selection step (Polzer *et al*. 2025). The strong selection in the first generation, where only the 18 highest-performing individuals (10 for R1 and 8 for R2) were mated, introduced a strong bottleneck in combination with efficient directional selection. While the resulting increase in linked selection contributed to the loss of allelic diversity (Charlesworth 2009), large effect loci contributing to traits under selection were likely fixed already within the first generation. Allele frequencies of loci with milder effects on trait expression also changed the strongest in the first generation (Figure 3c). This is in accordance with a simulation study by Pook *et al*. (2025) of the exact same experimental design indicating that the genetic gain reaches a plateau within a few generations if only the founding population is phenotyped and further selection is based solely on genomic information. Thus, the combination of drift and selection on major loci likely fixes highly beneficial alleles rapidly and depletes beneficial variation early on resulting in decreased efficacy of genome-based selection in consecutive generations, as the large effect loci might have dominated training of the prediction model. This is also supported by previous findings showing that retraining in each generation improves selection accuracy in later generations (Polzer *et al*. 2025). Although the genome-wide genetic diversity declined under breeding-imposed selection pressures over time, we were able to detect candidate loci, where beneficial alleles increased in frequency beyond the genome-wide allele frequency changes. Loci that showed the strongest signals of selection also changed their allele frequencies rapidly and were enriched for relevant gene ontology categories (Barghi *et al*. 2020). In addition, the selected loci significantly overlapped with QTL associated with the traits under selection in the experiment, namely, early vigor and plant height (Mayer *et al*. 2020) (Figure 4). Such an enrichment of potentially functional large-effect alleles has been previously associated with positive selection acting on standing genetic variation in model organisms and natural populations (Schlötterer 2023; Barrett and Schluter 2008). Here, we applied the principles of experimental evolution to a rapid plant breeding population and could show the genomic composition of selection gain over time. We were able to identify significant overlap of beneficial loci between replicates, suggesting the potential role of these loci in trait expression. Yet, most of the selected loci did not overlap between replicates, highlighting the likely role of polygenic inheritance of traits when selecting from standing genetic variation, even when applying intense selection in a short time frame.

In addition to the overlap of the positions of selection signals and QTL for traits under selection in the experiment, we detected signals of selection for traits, such as lodging and tillering, that were not under direct selection during the experiment. The selection on these traits might be attributed to unconscious selection in the field or correlated response through the pleiotropic control of traits (Hölker et al. 2019, Chen and Lübberstedt 2010; Gardner and Latta 2007). Such patterns have also been observed in tropical maize and model organisms, in which additional and sometimes unexpected traits have shown signs of being under selection (Choquette *et al*. 2023; Pfenninger and Foucault 2020; Teotonio *et al*. 2017; Burke and Rose 2009). These signals show that even highly controlled intentional selection for specific traits relies on fitness parameters beyond the individual traits under selection as individuals need to survive and complete their lifecycle to contribute to the consecutive generation.

Our results show that the population genetic signal of consecutive generations in a breeding program enables the identification of genomic regions that control traits under selection. However, intense selection rapidly reduces diversity. The regular introduction of genetic material, such as from landraces (Takou *et al*. 2025; Arca *et al*. 2023), could help supplement the genetic pool if they harbor novel beneficial alleles. European maize landraces are a rich source for such beneficial variation that can introduce novel alleles into breeding populations (Mayer *et al* 2020; Urzinger *et al* 2025). Genetic diversity in landraces is often underused, but using techniques like rapid cycling has the potential to harness this diversity to create high-yielding varieties that are both climate-adaptable and productive (Polzer *et al*. 2025, Rivera-Poulsen et al. 2026). In fact, simulation work showed that introducing novel diversity can balance short-term genetic gains and long-term diversity preservation, which is important for future breeding progress (Pook *et al*. 2025).

In this study, we showed that experimental evolution is a powerful approach for understanding how genetic diversity changes over time and how selection changes the allele frequencies of hundreds of loci simultaneously, but favors large effect loci in the short term. Plant breeding provides excellent populations to study this evolutionary question, as breeding programs are consecutive time-series of many generations. The combination of breeding populations and evolutionary genomics allow to pinpoint causal genetic loci controlling key traits in crops. Such systems will help us understand rapid evolution and can help improve crops in the future.

## Acknowledgments

We acknowledge KWS SAAT SE & Co. KGaA for development and preparation of plant material, genotyping, field experiments and data collection as part of the MAZE project. We thank A. Singh, Y. Zhang and T. Ali for fruitful discussions on analysis of time series datasets. The study was funded by the Federal Ministry of Research, Technology and Space (BMFTR, Germany) within the scope of the funding initiative “Plant Breeding Research for the Bioeconomy” (Funding ID: 031B1301A and 031B1301C) as part of the project MAZE 3.

## Author Contribution Statement

MGS and CCS conceived and designed the study. SdS prepared plant material and genotyping of the maize populations. MT and MTP conducted the analysis and prepared the figures. MT and MTP prepared the first draft of the manuscript. MGS, CCS and SdS read and provided feedback on the analysis. CCS and MGS edited the manuscript. All authors read and approved the manuscript.

## Data archiving

All data and material are available as described in Polzer et al., 2025.

## Research Ethics Statement

Not applicable.

## Supplement

**Figure S1.**
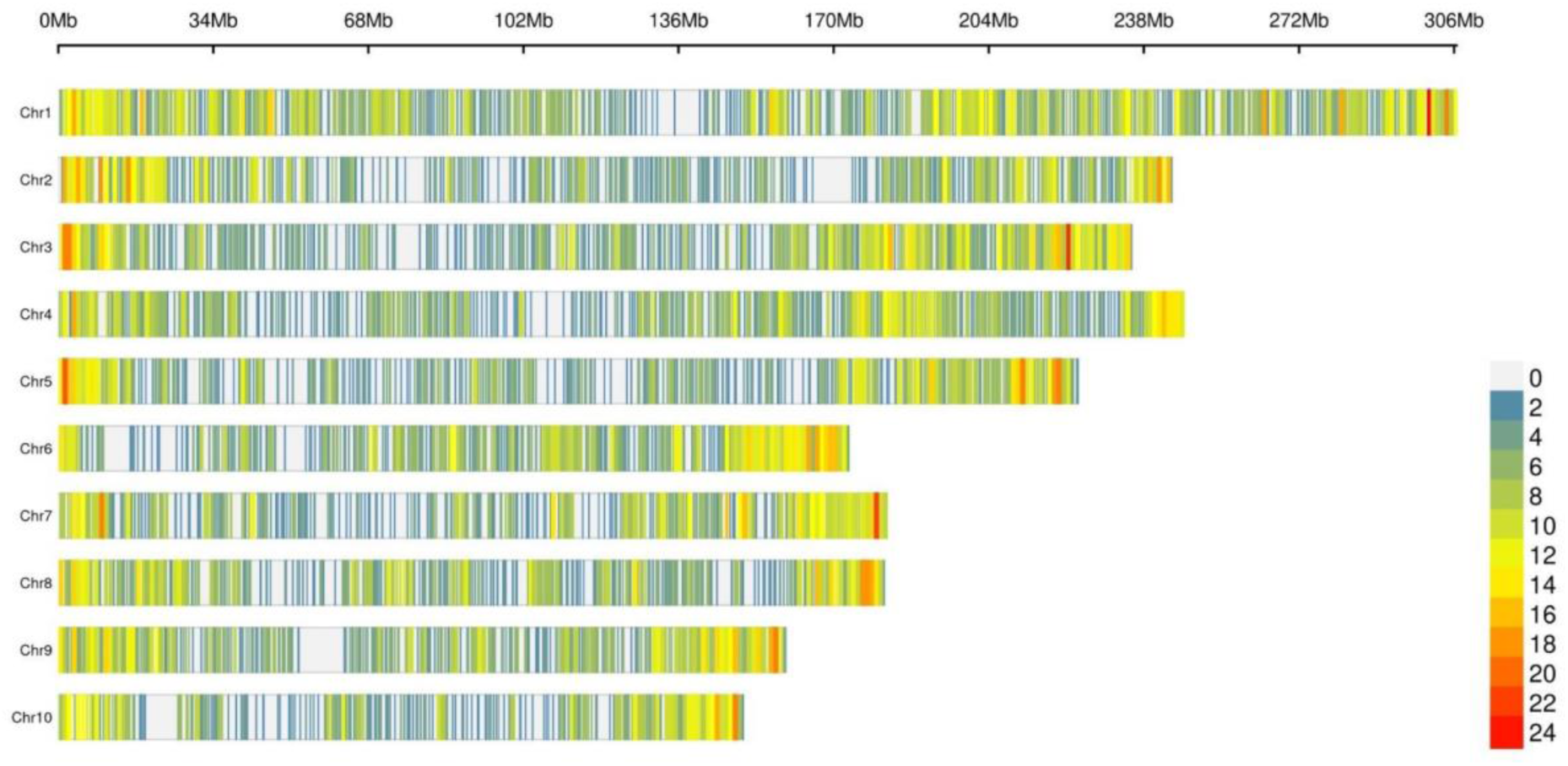
Distribution of the 11,160 SNP dataset. Genome-wide density across chromosomes calculated in 1 Mb windows.

**Figure S2.**
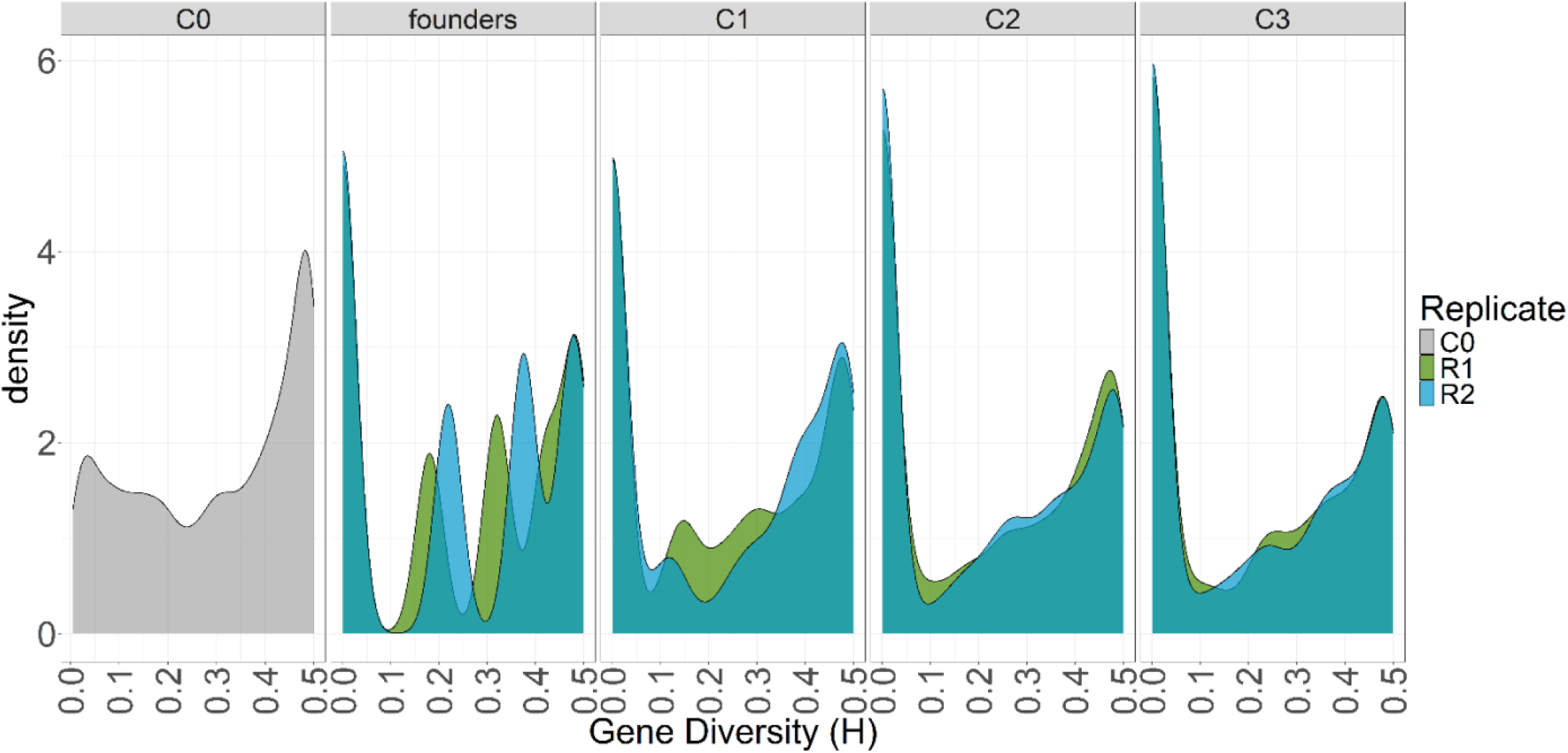
Gene diversity (H) density distributions from the base population C0, founder lines, and subsequent populations C1–C3.

**Figure S3.**
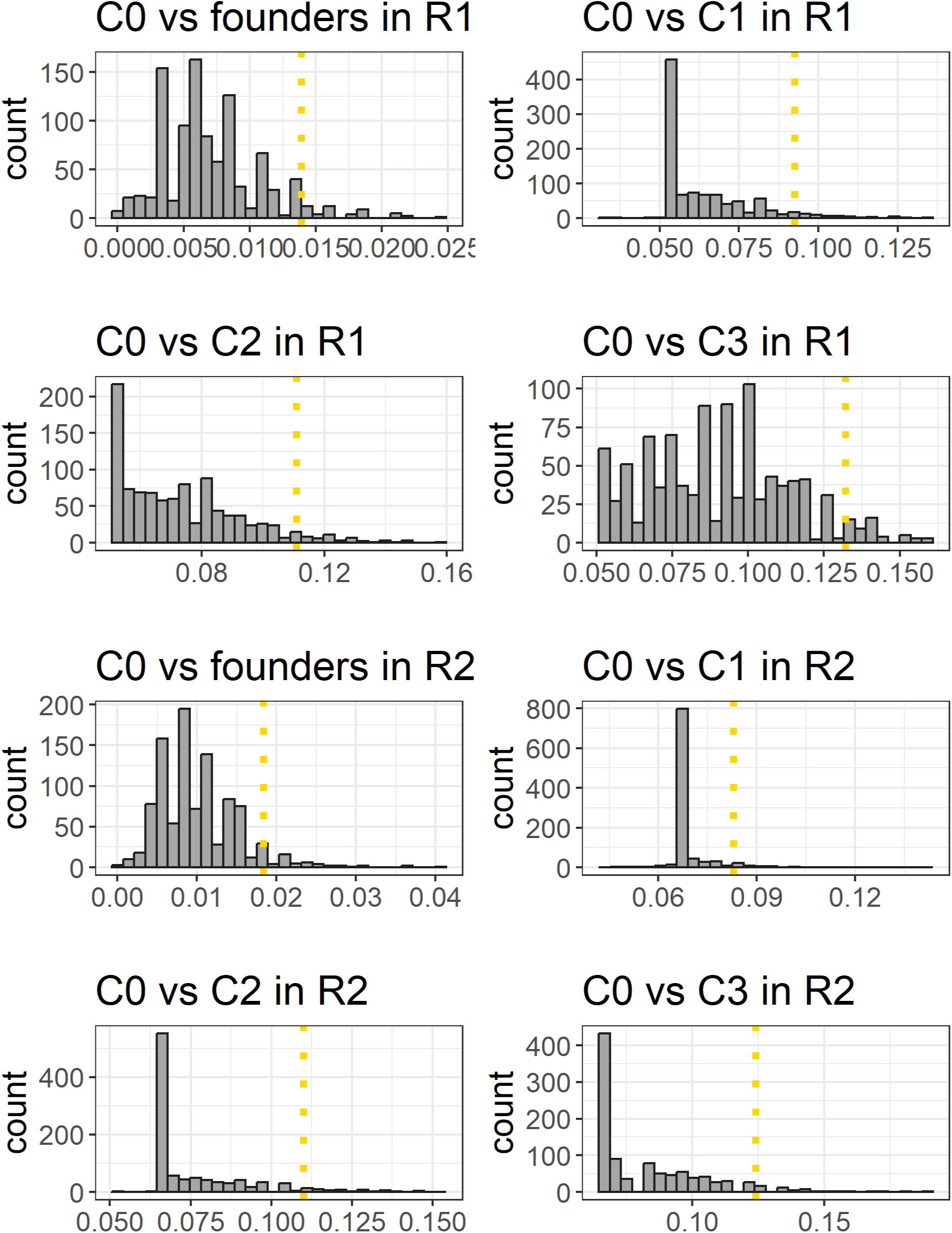
The distributions of the *F_st_* values in the permutation datasets, where the sample identity was randomized. The *F_st_* were estimated between each population to C0 for both replicates. The 99^th^ percentile of each distribution is marked with the dotted lines.

**Figure S4.**
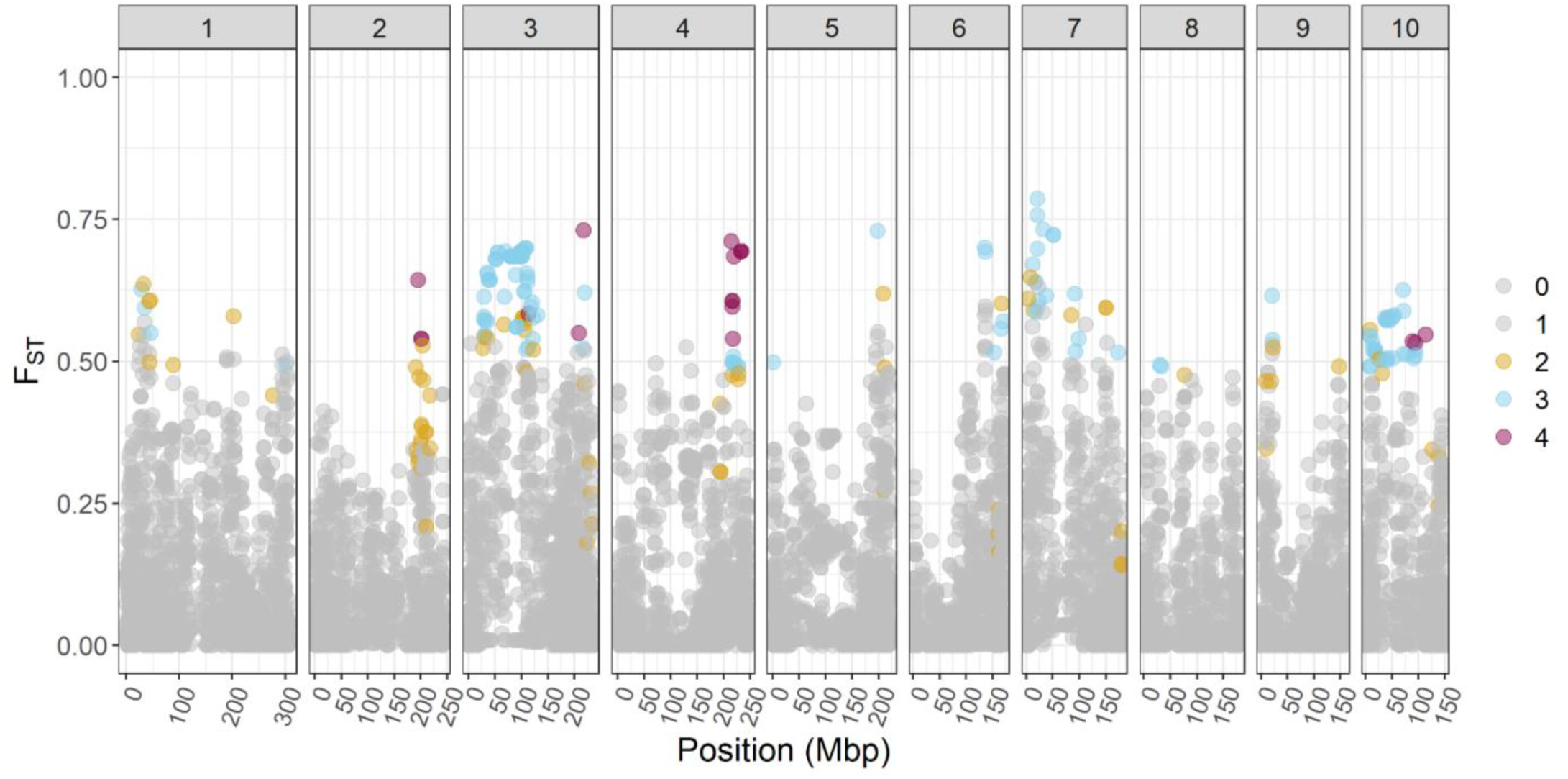
The *F_ST_* values between C0 and C3 indicate changes in R2. The colored dots in both plots indicate in how many of the comparisons between C0 and founder lines, C1, C2, and C3, the locus was an outlier for *F_ST_*. Each facet corresponds to one chromosome.

